# Mutation of two intronic nucleotides alters RNA structure and dynamics inhibiting MBNL1 and RBFOX1 regulated splicing of the Insulin Receptor

**DOI:** 10.1101/2024.01.08.574689

**Authors:** Zohreh R. Nowzari, Melissa Hale, Joseph Ellis, Samantha Biaesch, Sweta Vangaveti, Kaalak Reddy, Alan A. Chen, J. Andrew Berglund

**Affiliations:** Department of Chemistry, University at Albany–SUNY, Albany, NY 12222, United States; The RNA Institute, University at Albany–SUNY, Albany, NY 12222, United States; Department of Neurology, School of Medicine, Virginia Commonwealth University, Richmond, Virginia 23298, United States; Department of Biochemistry & Molecular Biology, Center for NeuroGenetics, College of Medicine, University of Florida, Gainesville, Florida 32610, United States; Department of Biology, University at Albany–SUNY, Albany, NY 12222, United States

## Abstract

Alternative splicing (AS) of Exon 11 of the Insulin Receptor (*INSR*) is highly regulated and disrupted in several human disorders. To better understand *INSR* exon 11 AS regulation, splicing activity of an *INSR* exon 11 minigene reporter was measured across a gradient of the AS regulator muscleblind-like 1 protein (MBNL1). The RNA-binding protein Fox-1 (RBFOX1) was added to determine its impact on MBNL1-regulated splicing. The role of the RBFOX1 UGCAUG binding site within intron 11 was assessed across the MBNL1 gradient. Mutating the UGCAUG motif inhibited RBFOX1 regulation of exon 11 and had the unexpected effect of reducing MBNL1 regulation of this exon. Molecular dynamics simulations showed that exon 11 and the adjacent RNA adopts a dynamically stable conformation. Mutation of the RBFOX1 binding site altered RNA structure and dynamics, while a mutation that created an optimal MBNL1 binding site at the RBFOX1 site shifted the RNA back to wild type. An antisense oligonucleotide (ASO) was used to confirm the structure in this region of the pre-mRNA. This example of intronic mutations shifting pre-mRNA structure and dynamics to modulate splicing suggests RNA structure and dynamics should be taken into consideration for AS regulation and therapeutic interventions targeting pre-mRNA.

**Abstract figure:**
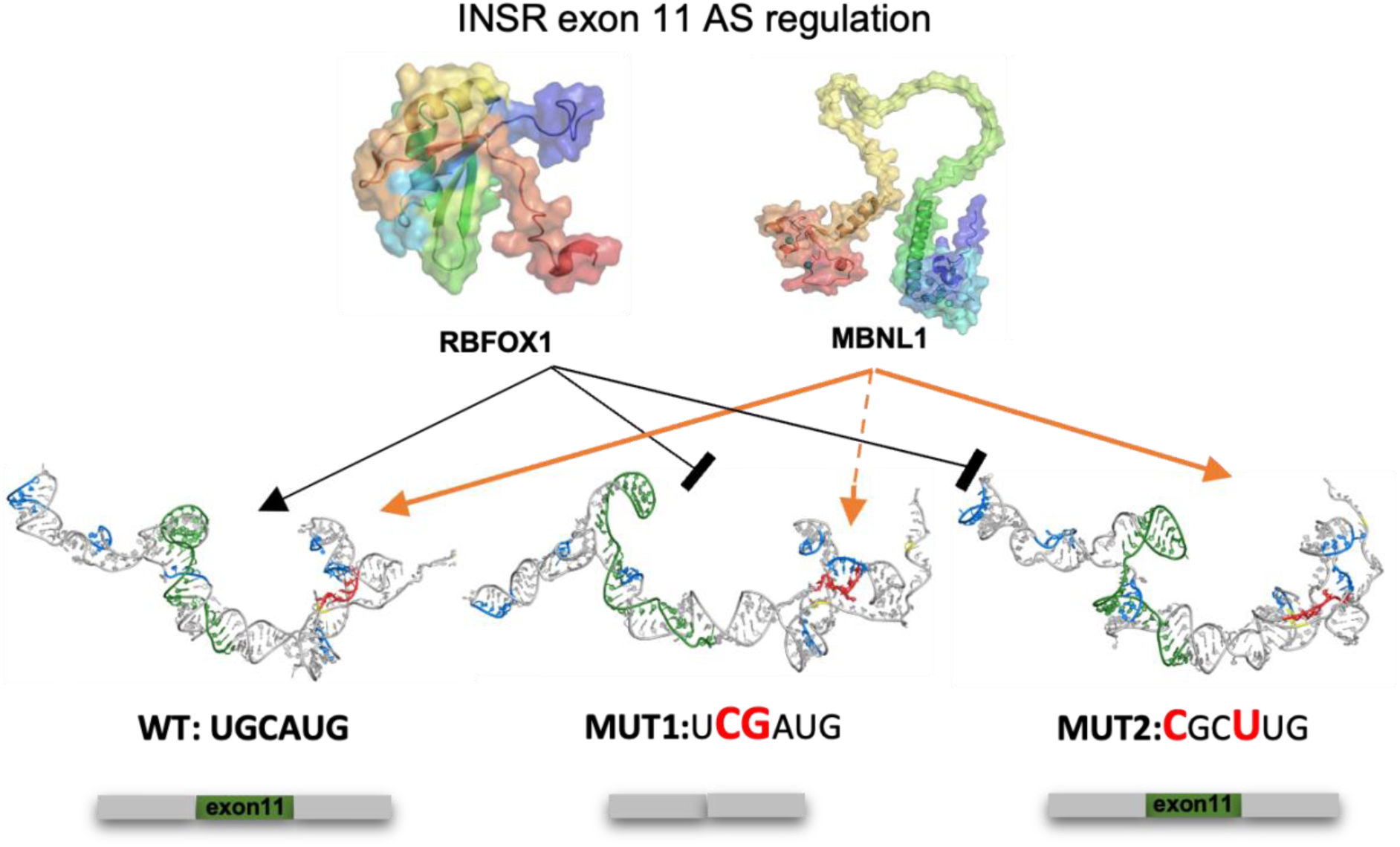
Model for *INSR* exon 11 splicing regulation through the UGCAUG motif. We propose that the UGCAUG motif, MBNL1, and RBFOX1 cooperatively regulate *INSR* exon 11 splicing. Mutating this UGCAUG motif is sufficient to alter RNA structural dynamics to disrupt this regulation.

## Introduction

Alternative splicing (AS) is a crucial step in gene expression by which a single gene can produce multiple distinct transcripts, thus diversifying the transcriptome and proteome. AS is regulated by a complex network of pre-mRNA regulatory sequence elements and trans-acting protein factors (1–4). While the mechanism of AS has been well studied, there are unanswered questions remaining. For example, how multiple RNA binding proteins (RBPs) function cooperatively or antagonistically through pre-mRNA sequence elements and their structural configurations to regulate splicing is an open and important question in the field. RBPs are a class of proteins with essential roles in gene regulation, one of which is their role in regulating AS. The Muscleblind-like (MBNL) family of RBPs is one such group of AS regulators with important roles in development and pathogenesis of Myotonic Dystrophy (DM) (2, 5–11).

MBNL proteins have been shown to bind to YGCY cis-regulatory motifs to regulate AS outcomes of their target pre-mRNAs in a positional-dependent manner (12, 13). Specifically, MBNL proteins generally promote cassette exon inclusion by binding to downstream RNA motifs and repressing or blocking exon inclusion by binding upstream intronic or exonic elements (14–17). In addition to MBNL proteins, other RBPs such as NOVA, RBFOX, and CELF exhibit positional-dependent patterns in regulating AS outcomes (18–26).

Recent work has demonstrated an overlap in AS regulation by MBNL and RBFOX proteins in skeletal muscle, and this cooperative interaction has been proposed to drive transcriptional changes significant for stem cell differentiation (27, 28). Previous reports identified an MBNL response element within intron 11 of the *INSR* gene (downstream of exon 11) critical for proper regulation by MBNL proteins (14, 29). Knockdowns of MBNL1 and RBFOX1 showed changes in *INSR* splicing in fibroblasts (27), and most recently, RBFOX2 was shown to mediate INSR exon 11 inclusion (30).

The *INSR* gene encodes for the insulin receptor (IR), which is critical in the insulin signalling pathway in regulating glucose uptake and release (31). Mutations in the *INSR* gene have been linked to severe insulin resistance, including leprechaunism, Rabson-Mendenhall syndrome, or Insulin-resistance syndrome type A (32–34). AS of *INSR* is cell type-specific, and the relative proportions of the different isoforms vary during development, aging, and disease states (35). AS of *INSR* exon 11, which generates two isoforms that differ by 12 amino acids in the hormone binding domain (36, 37), is a well-studied AS event. In humans, both the exclusion (IR-A) and inclusion (IR-B) isoforms of exon 11 are present in tissues such as skeletal muscle, adipose tissue, kidney, placenta, and heart; however, IR-B is the predominant isoform found in insulin-responsive tissues (35). Furthermore, aberrant AS of IR exon 11 is found in Type II Diabetes, Cancer, and Myotonic Dystrophy types I and II (DM1 and DM2) (38–41).

In this study, to better understand the sequence, structural, and protein interactions governing AS outcomes, we adopted a multifaceted and integrated approach focusing on *INSR* exon 11 as a model AS inclusion event using cellular assays and computational modelling. Using an *INSR* minigene reporter system (42), we tested the activity of a single UGCAUG motif in *INSR* intron 11 that had previously been shown to be an important control element for AS regulation of exon 11 (30). We performed mutational analysis and splicing studies that suggest that MBNL1 and RBFOX1 proteins both work through this UGCAUG motif to regulate AS of *INSR* exon 11. All-atom molecular dynamics simulations (MDS) revealed that a region of the *INSR* pre-mRNA adopts a stable RNA structure disrupted by two point mutations in the UGCAUG motif, which can be rescued by two additional mutations. An ASO was used to test the importance of a predicted helix, and treatment with this ASO altered AS regulation of *INSR* exon 11 consistent with predicted RNA structure and dynamics. The results indicate the importance of considering RNA structure and dynamics in the analysis of AS regulation and suggest that a highly structured RNA interacting with multiple RBPs plays an important role in regulating *INSR* exon 11 AS.

## Materials and Method

### Plasmids and cloning

N-terminal HA-tagged RBFOX1 (1-397 amino acids, RNA binding protein fox-1 homolog isoform 4; NCBI accession number NP_061193.2) was synthesized (GenScript, Piscataway, NJ, USA) and cloned into pcDNA3.1+ (Life Technologies, Carlsbad, CA, USA) using standard cloning procedures. Mutations to the wild type *INSR* minigene (36) were performed using the Quick Change Site-Directed Mutagenesis platform (Agilent Technologies, Santa Clara, CA, USA) and Phusion high-fidelity DNA polymerase (New England Biolabs (NEB), Ipswich, MA, USA).

### Antisense oligonucleotides

2’-O-methyl-based ASOs were ordered from IDT (Integrated DNA Technologies, Coralville, IA, USA). The 3’ splice site (3’SS) blocker ASO is {5’-AGAGGUUUUUCUGUGGAAAC-3’(L=20nt)}, 5’ splice site (5’SS) ASO is {5’-GGUGAGUCAUACCUAGGGUC-3’(L=20nt) and 3’ compelmentry ASO (3’COMP) is {5’-AACCCCUGGGUUCUCCGA-3’(L=18nt)}. All ASOs were resuspended in nuclease-free water to a final concentration of 100 μM. The ASOs were transfected into MBNL1/RBFOX1 dual inducible HEK-293 cell line (ddHEK) (17) using Lipofectamine 3000 (Thermofisher, Waltham, MA, USA) according to the manufacturer’s instructions.

### Cell culture and transfection

HA-MBNL1 and ddHEK were routinely cultured as a monolayer in Dulbecco’s modified Eagle’s medium (DMEM) (Corning, Corning, NY, USA) supplemented with 10 % fetal bovine serum (FBS, Corning), 10 μg/mL blasticidin (Gibco), and 150 μg/mL hygromycin (Gibco) at 37 °C under 5 % CO_2_. HA-MBNL1 cells were plated in twenty-four well plates at a 1.5 x 10^5^ cells/well density for all minigene work. Cells were transfected 24 hours later at approximately 80% confluence. Plasmids were transfected into each well with 1.5 μl of TransIT-293 (Mirus Bio) per the manufacturer’s protocol, 250 ng of empty vector (pcDNA3.1+, mock) or the pcDNA3.1+ HA-RBFOX1 expression vector was co-transfected with 250ng of a single minigene reporter (total = 500 ng/well). Fresh Doxycycline (Sigma Aldrich) was prepared at 1 mg/mL, diluted, and added to the cells at the appropriate concentrations (0.16, 0.31, 0.62, 1.2, 2.5, 5.0, 10 ng/mL) four hours post-transfection. Twenty-four hours post-transfection, cells were harvested for experimental analysis. For ASO transfections, ddHEK cells were plated at a density of 2×10^5^ cells/well in a 12-well plate. Cells were allowed to adhere overnight and then transfected with the appropriate ASO at a final concentration of 25 nM. 10ng/mL fresh Doxycycline (Sigma Aldrich) was added simultaneously to induce MBNL1 expression with the ASO and transfection reagent. Cells were harvested at 48 h post-transfection.

### Cell-based splicing assays and curve fitting

Total RNA from HA-MBNL1 and ddHEK cells was isolated using the RNeasy Mini kit (Qiagen, Germantown, MD, USA). The isolated RNA was processed via reverse-transcription (RT)-PCR, and PSI (percent spliced in) for each minigene event was calculated as previously described (17). 1000 ng of RNA was reverse-transcribed (or 500 ng for the ASO protocol) using SuperScript IV (Invitrogen, Carlsbad, CA, USA) with random hexamer priming (Integrated DNA Technologies, Coralville, IA, USA) according to the manufacturer’s protocol, except that half of the recommended SuperScript IV enzyme was utilized. cDNA was then PCR amplified for 25-32 cycles using flanking exon-specific primers (forward primer = CAACCAGAGTGAGTATGAGGATT and reverse primer = CCGTCACATTCCCAACATCG, annealing temperatures = 55). PCR-amplified cDNA samples were visualized and quantified using the Fragment Analyzer DNF-905 dsDNA 905 reagent kit, 1-500bp (Agilent Technologies), and associated ProSize data analysis software. Quantified PSI values were plotted against relative MBNL levels determined by immunoblot or log (Doxycycline (ng/mL)). Curve fitting was performed using the GraphPad Prism software using non-linear curve fitting (log(agonist) vs. response – variable slope (four parameters)) (PSI = PSI_min_+ ((PSI_max_-PSI_min_) / (1 + 10^((log(EC50) − log[MBNL1])^ * ^slope)^). Where appropriate, the minimum and maximum values were restricted to fall between 0 and 1, respectively. Parameters correlating to biological data, such as log (EC50) and slope, were derived from these curves (n=3).

### RNA structure folding and dynamics

RNA Composer (43), a computational tool for RNA structure prediction, was used to generate the initial three-dimensional (3D) structure of the 249 RNA fragment that contains 52 nucleotides of intron 10, including two mapped branch sites (42), 36 nucleotides of exon 11, and 161 nucleotides of intron 11 that contains one UGCAUG RBFOX1 site and six YGCY MBNL1 binding sites (M-BS). From the ensemble of initial structures, we selected the centroid structure as the representative model (44) and performed structure refinement steps, including energy minimization and equilibration (17 nanoseconds (ns)). The mutations in the RBFOX1 motif (MUT1(U<u>CG</u>AUG)) and MUT2(<u>C</u>GC<u>U</u>UG)) were introduced into the generated 3D structure. Next, three independent 90 ns molecular dynamics (MD) simulations were run, using GROMACS (45), for wild type (WT), MUT1, and MUT2 structures. The Amber-99-Chen-Garcia force field with backbone phosphate modifications was employed for nucleic acids (46). In the simulation setup, a cubic box with dimensions of 22.74 nm on each side was created to accommodate the pre-mRNA structure and the surrounding environment, 1, 558, 595 atoms in total. The system was simulated in a 0.1M KCl solution, which included 1, 300 K+ ions, 1, 052 Cl-ions (47), and 387, 063 TIP4P-Ewald water molecules (48). The time step used was 2 femtoseconds (fs), allowing us to capture the molecular motions over time accurately. The system was maintained at a constant temperature of 300 Kelvin (K) using the V-rescale thermostat (49). For pressure control, we utilized the Berendsen barostat with a reference pressure of 1 bar, maintaining isotropic pressure scaling (49).

## Results and Discussion

### Characterization of MBNL1-dependent alternative splicing of *INSR* exon 11 minigene reporter

A previously published minigene reporter system (42) was used to evaluate RNA sequence elements that control MBNL1 and RBFOX1 regulation of *INSR* exon 11 AS (Figure 1). *INSR* exon 11 splicing has been extensively studied, with several MBNL consensus YGCY motifs identified as critical for MBNL-dependent exon 11 inclusion (14, 29) and a UGCAUG RBFOX binding site shown to be necessary for regulation of *INSR* exon 11 by RBFOX2 (Figure 1). Of note, there is a second RBFOX GCAUG binding motif ∼180nt downstream from exon 11, but it was shown to not be essential for proper splicing of *INSR* exon 11 (30).

**Figure 1.**
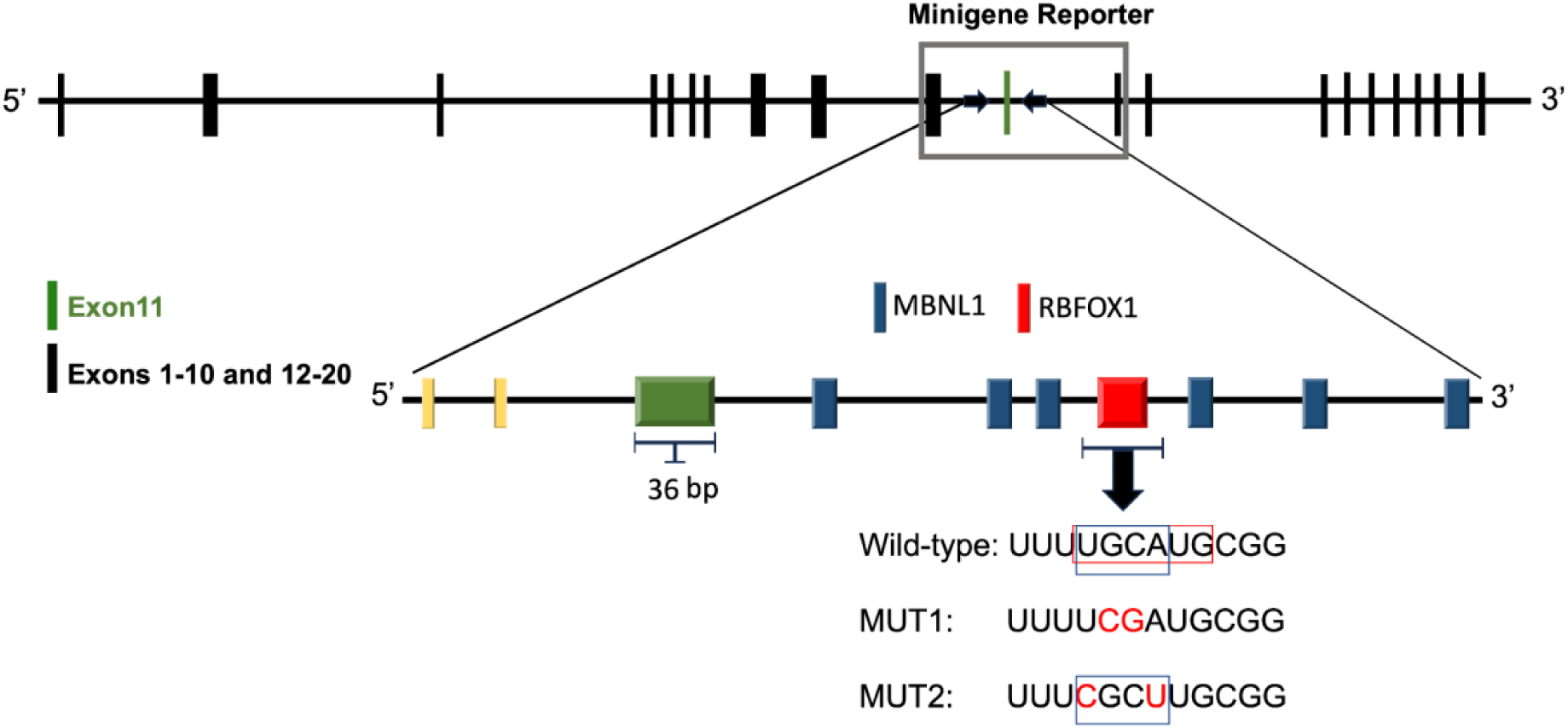
The figure illustrates the *INSR*, which spans 182, 160 nucleotides (nt). Exons are represented by black boxes indicating exons 1 to 10 and 12 to 22, and a green box representing exon 11. The minigene region of interest, encompassing exon 10 to exon 12, is highlighted by a grey box. The bottom part of the schematic depicts the presence of recognition sites for two important regulatory proteins, MBNL1 and RBFOX1; this segment of the gene was the substrate for the in-silico modelling (249 nt). Moreover, the nucleotide sequences of R-BS are provided in the last part. The first line corresponds to the WT sequence, while the subsequent two lines depict the sequences of the two mutant variants, MUT1 and MUT2 (the mutated nucleotides shown in red).

We first compared minigene and endogenous *INSR* exon 11 splicing in a tetracycline-inducible HA-MBNL1 HEK-293 cell line in which *INSR* exon 11 AS responds to a dose range of MBNL1 protein (50). The *INSR* minigene reporter was transfected, and MBNL1 was induced across an 11-point dose range of doxycycline. Percent spliced in (PSI) of *INSR* exon 11 was measured and plotted as a function of the log of the concentration of Doxycycline (i.e., log([Dox])), a proxy for HA-MBNL1 protein concentration (Figure 2A, blue curve). A four-parameter dose-response curve can then be fit to derive quantitative measures that describe the observed patterns of AS regulation, such as EC50 (amount of MBNL1 protein required to obtain splicing regulation at 50% of maximum PSI), slope (relative measure of MBNL1 cooperativity), and deltaPSI (difference in PSI between the top and bottom of the dose-response curves) (50). We found that the minigene had a more modest deltaPSI value of 0.31 (Figure 2A blue curve, D) compared to the endogenous *INSR* exon 11 PSI value of ∼0.6 (50). Furthermore, shifts in slope (2.3 vs. 1.1) and LogEC50 (0.23 vs. −0.07) suggested that MBNL1 AS regulation was more cooperative in the minigene system versus the endogenous pre-mRNA and that the minigene *INSR* mRNA required a higher concentration of MBNL1 protein to regulate splicing. These results are consistent with a greater requirement of MBNL1 protein to regulate the plasmid-derived exogenous minigene RNA, which is expected to be at a higher concentration than endogenous *INSR* pre-mRNA. It should also be noted that the *INSR* minigene lacks part of intron 11 (a sizeable internal deletion leaving ∼180 nucleotides at both the 5′ and 3′ ends), which may affect AS regulation. However, the conservation of AS regulation by MBNL1 in the exogenous and endogenous *INSR* exon 11 pre-mRNA validated the use of the *INSR* minigene system as a tractable model to assess the regulatory mechanisms of *INSR* exon 11 AS.

**Figure 2.**
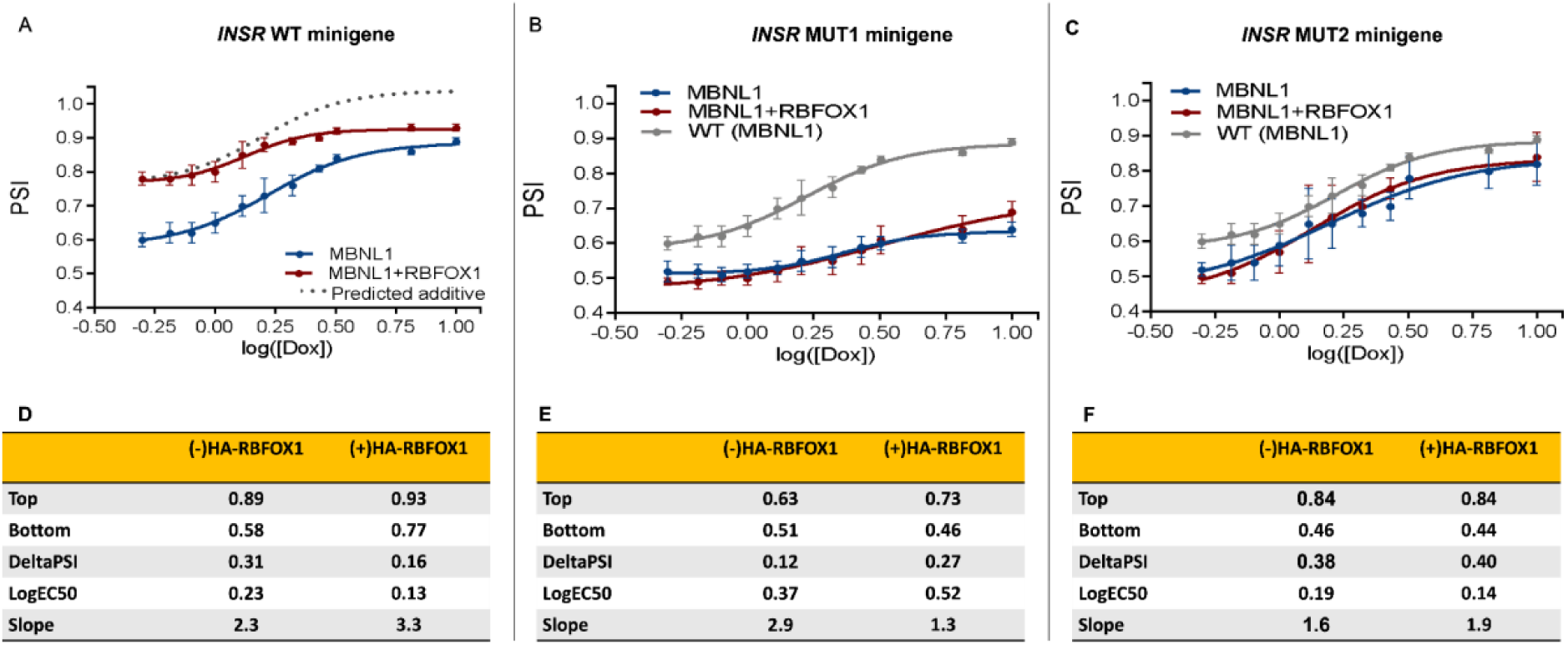
RBFOX1 Modulation of MBNL1-Dependent *INSR* Exon 11 Splicing Regulation; Plots (**A-C**) show changes in *INSR* exon 11 inclusion levels across a gradient of MBNL1 protein expression, with and without HA-RBFOX1 co-transfection in WT, MUT1 and MUT2 respectively. **(D-F)** Quantitative values correspond to the plot above, including deltaPSI, LogEC50, and slope.

### RBFOX1 expression modulates MBNL1 dose-dependent splicing regulation of wild type *INSR* minigene splicing reporter

To determine if RBFOX1 modulates MBNL1-dependent splicing of *INSR* exon 11, an HA-RBFOX1 expression vector was co-transfected with the minigene reporter, and MBNL1 was induced using doxycycline as described above. *INSR* exon 11 PSI was measured in the presence and absence of transfected RBFOX1, and dose-response curves were plotted. Co-expression of HA-RBFOX1 led to an overall reduction in deltaPSI across the MBNL1 concentration gradient compared to MBNL1 expression alone (0.16 vs 0.31, Figure 2A, D), demonstrating that RBFOX1 modulates MBNL1 regulation of *INSR* exon 11 AS in this system.

Interestingly, the observed change in PSI occurred non-uniformly across the MBNL1 dose gradient. While the lowest levels of MBNL1 with co-expression of HA-RBFOX1 resulted in a 0.19 increase in PSI of *INSR* exon 11 inclusion (0.58 vs 0.77, Figure 2D; Bottom), induction of higher cellular concentrations of MBNL1 resulted in only a 0.04 minimal increase in PSI (0.89 vs 0.93, Figure 2D; Top). Based on the deltaPSI value at the lowest MBNL1 induction, a uniform additive effect would be predicted to exceed the maximum level of exon inclusion (PSI = 1.0) detectable in our assay (Figure 2A, dotted Curve). Thus, the non-uniform deltaPSI resulting from the co-expression of RBFOX1 may reflect the saturation point of the *INSR* minigene splicing event. However, if MBNL1 and RBFOX1 regulated this exon in a genuinely independent manner, we expect the EC50 values to be similar with and without the expression of HA-RBFOX1. This is not the case, but rather, less MBNL1 is required for an equivalent amount of splicing regulation when HA-RBFOX1 is co-expressed (0.22 vs. 0.13; Figure 2D).

These dosing studies indicate that the observed pattern of AS regulation could be occurring through a cooperative mechanism involving both RBPs. This is further supported by the observed change in slope value, albeit modest, with and without co-expression of RBFOX1 (3.3 vs. 2.3, respectively; Figure 2D). While previous work has demonstrated MBNL and RBFOX coordinate splicing decisions, there is limited mechanistic evidence demonstrating how these two proteins modulate AS of the same pre-mRNA targets.

### Mutagenesis revealed a shared RNA cis-regulatory element through which MBNL1 and RBFOX1 co-regulate *INSR* exon 11 inclusion

To further dissect the mechanism through which RBFOX1 modulates the MBNL1 dose-response curve of *INSR* exon 11 inclusion, we chose first to reconfirm the importance of the UGCAUG predicted RBFOX binding site within the downstream intron 11. Nakura and colleagues (30) used RBFOX2 to demonstrate that the UGCAUG motif is critical for regulating *INSR* exon 11. Because the RNA recognition motif (RRM) is identical for RBFOX1 and RBFOX2 (19), we hypothesize that this lone site is critical for RBFOX1 or RBFOX2 to be able to modulate MBNL1-dependent splicing of *INSR* exon 11 effectively. We mutated the lone UGCAUG motif to U<u>CG</u>AUG to generate the *INSR* MUT1 minigene (Figure 2B), as previous work has shown that inversion of this GC dinucleotide abrogates RBFOX1 binding site(R-BS) and prevents AS regulation (18, 51, 52). Interestingly, without co-expression of RBFOX1, we found this mutation significantly blunted MBNL1 dose-dependent splicing regulation in *INSR* MUT1 (Figure 2B, compare grey and blue curves) compared to WT *INSR* (Figure 2A). As expected, though, mutation of this site nearly eliminated the effect of transfected RBFOX1 on AS of *INSR* exon 11 (Figure 2B, red curve). This result is consistent with this RBFOX binding site being critical for regulating *INSR* exon 11 inclusion by RBFOX1/2. The LogEC50s of MUT1, both in the absence and presence of co-expressed RBFOX1, increased compared to the WT values. The co-expression of RBFOX1 further increased the LogEC50 for MUT1 compared to the MBNL only dose-response curve (0.52 vs. 0.37; Figure 2E). Interestingly, there was also a decrease in slope (2.9 vs 1.3, Figure 2E) with co-expression of RBFOX1. These results indicate the U<u>CG</u>AUG mutation weakens MBNL1 cooperativity, and more MBNL1 protein is required to achieve reduced AS regulation.

Overall, the substantial impact of MUT1 on MBNL1-dependent splicing regulation was unexpected given that (1) there are six M-BS present within intron 11 that have previously been shown to facilitate MBNL1-dependent exon 11 inclusion (14, 15, 29, 42), and (2) UGCAUG does not contain what is considered an MBNL1 high-affinity motif (12), but rather a UGCA which has been identified as a lower affinity M-BS (53). Thus, we predict MBNL would prefer to bind the numerous other consensus MBNL1 motifs surrounding this site before the less preferred UGCA. By changing the motif to <u>C</u>GC<u>U</u>UG (MUT2), we created a YGCY motif, which is not predicted to be bound by RBFOX1. As predicted, the co-expression of HA-RBFOX1 did not modulate AS of *INSR* exon 11 MUT2 (Figure 2C). The introduction of the new YGCY motif significantly restored MBNL1’s ability to regulate the splicing of *INSR* exon 11 with activity more similar to WT levels in the absence of co-expressed RBFOX1 (Figure 2C). The cooperativity for MUT2 (1.6 and 1.9, Figure 2F, compared to 2.3 and 3.3, Figure 2D) is lower compared to WT, while the LogEC50 is similar to WT, suggesting that the newly introduced YGCY site allows splicing regulation at lower MBNL1 concentrations. Why the introduction of a seventh YGCY motif would have this impact led us to consider how MBNL1 and RBFOX1 interact with their binding sites.

### RNA folding and dynamics of the *INSR* exon 11 and MBNL1 and RBFOX1 binding sites

Significant work on AS regulation by MBNL proteins has shown that MBNL proteins bind single-stranded RNA and can stabilize stem-loop structures in their target pre-mRNA substrates (54–57). To explore if the RNA structure of the pre-mRNA could be altered by the studied mutations, we performed RNA structure folding and dynamics studies on a 249 fragment of the *INSR* pre-mRNA that contained 52 nucleotides of intron 10, 36 nucleotides of exon 11, and 161 nucleotides of intron 11 (Figure 3A). An all-atom molecular dynamics simulation (MDS) was performed on the 249 fragment of the *INSR* pre-mRNA (Figure 3B). These simulations allowed us to observe and analyze the dynamic behavior of individual atoms within the mRNA structures over time (58). Specifically, we focused on understanding the consequences of the experimentally introduced mutations at the RBFOX motif. The structure generation and simulation details are described in the methods. Briefly, we performed 90 ns all-atom MDS on three systems: WT, MUT1, and MUT2. We evaluated two structural indicators: solvent-accessible surface area (SASA) and root mean square fluctuation (RMSF), as described in the methods.

**Figure 3.**
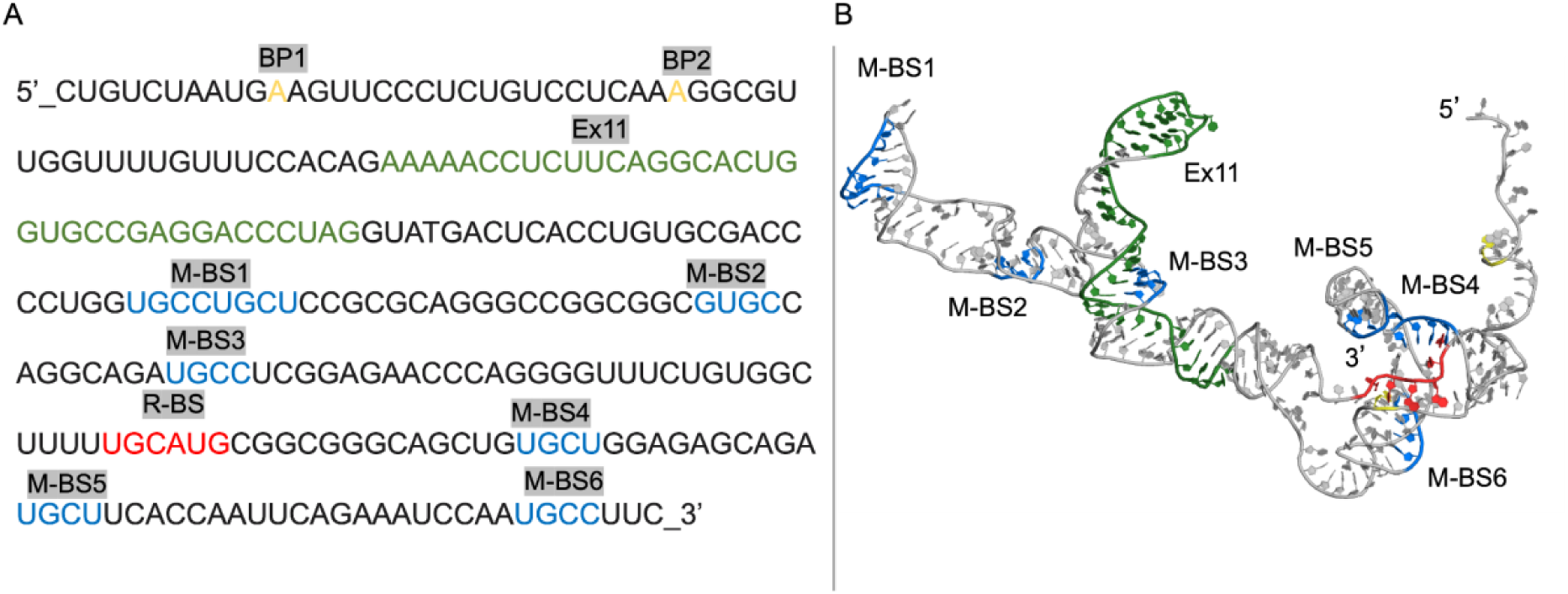
**(A)** Sequence of the 249 nucleotides of *INSR* exon 11 and adjacent sequences with mapped branchpoints in yellow (BP), exon 11 in green, MBNL binding sites in blue (M-BS), and RBFOX1 binding site in red (R-BS). **(B)** Modelled tertiary structure of the 249 nucleotide *INSR* pre-mRNA.

### Point mutations change solvent-accessibility and stability of the *INSR* pre-mRNA

SASA values are typically measured in square nanometers (nm^2^) per nucleotide, and they represent the surface area of each nucleotide that is exposed to solvent molecules during the simulation. SASA values indicate how much of a nucleotide’s surface is “visible” to the surrounding environment. A higher SASA value suggests that a nucleotide or region is more exposed or accessible. In contrast, a lower SASA value implies that it is more shielded from the surrounding solvent (59). In our study, we applied SASA to gauge MUT1 and MUT2 induced pre-mRNA conformational changes compared to the WT pre-mRNA structure.

The SASA analysis allowed us to assess changes in the accessibility of critical RNA motifs, including all MBNL1 recognition sites(M-BS), the 3’SS and 5’SS of exon 11, and the R-BS. Shown in (Figure 4 A-D) is the accessible surface area for exon 11, R-BS, the 5’, and the 3’SS in the MUT1, MUT2, and WT pre-mRNAs. We find that in MUT1, both exon 11 (Figure 4A) and the 5’SS (Figure 4B) are more solvent-exposed with an average SASA (nm^2^) score per nucleotide of 2.6 and 2.4 compared to WT at 2.3 and 2.1 and MUT2 at 2.4 and 1.9 respectively. The SASA score for the 3’SS is similar in all three cases, as evident from the overlapping values in Figure 4C. For R-BS, the SASA trends differ from the other motifs, with MUT2 having a more exposed site compared to WT and MUT1 (Figure 4D). The overall similarity in SASA scores for WT and MUT2 are consistent with these two pre-mRNAs being regulated by MBNL1 in a similar manner, while the changes in MUT1 may explain why MBNL1 has blunted splicing activity with this pre-mRNA substrate.

**Figure 4.**
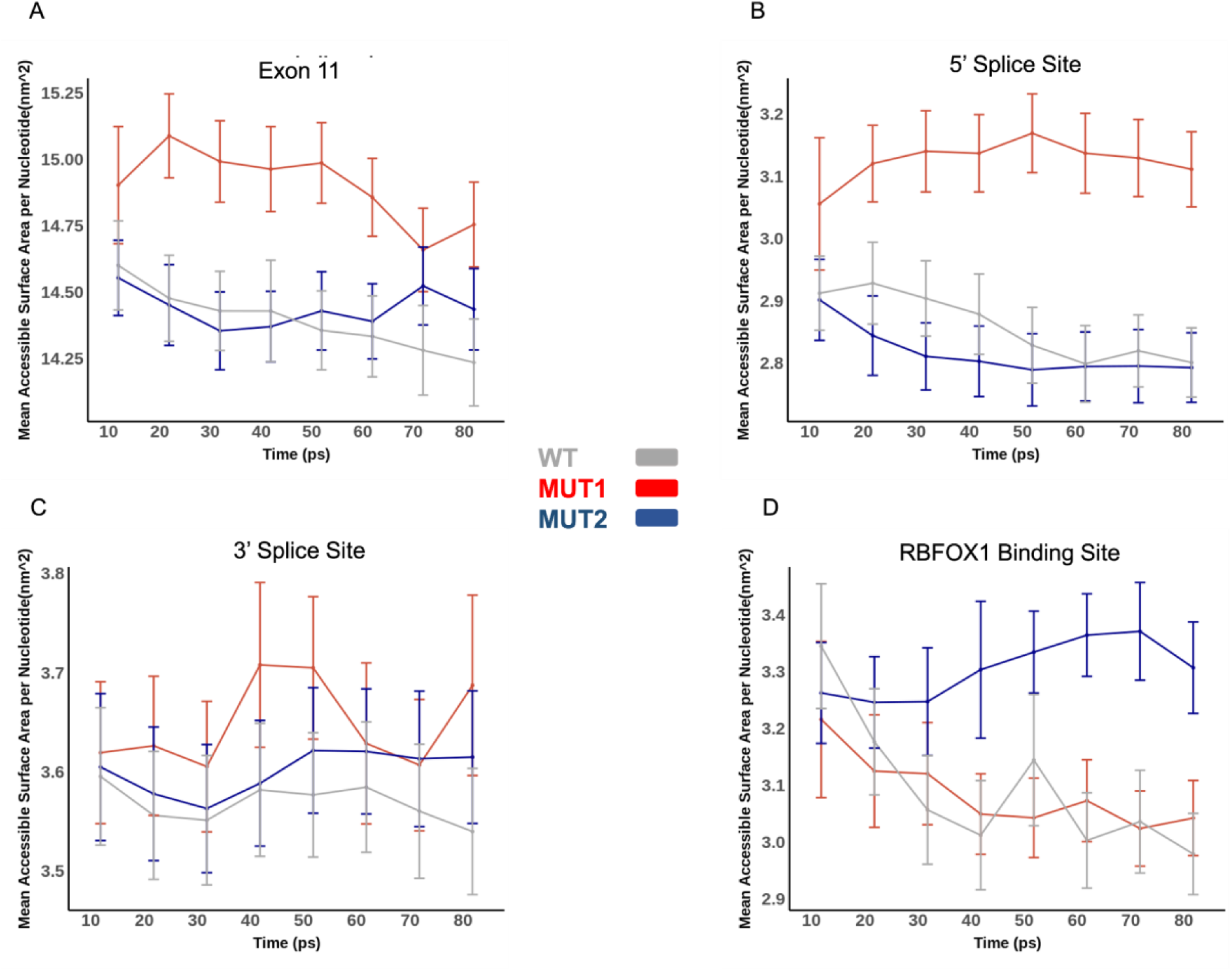
Graphs (**A, B, C, and D**) represent the solvent-accessible surface area (SASA) analysis, WT(gray), MUT1(red), and MUT2(blue), for different regions in the pre-mRNA structure: exon 11, the 3’SS and 5’SS, and the R-BS, respectively. The error bars represent one standard deviation (SD) above and below the mean calculated over every 5, 000 frames, measuring the structural variability during the simulations. On the x-axis are different simulation windows, each corresponding to specific time intervals (e.g., 5000 frames) or simulation stages during the 90 ns trajectory. Eight windows on the x-axis indicate various points in the simulation trajectory, allowing us to study the structural changes over time. The y-axis represents the SASA of the pre-mRNA region, which provides insights into the exposed surface area of the RNA and its interactions with surrounding molecules.

To better understand how the SASA trends manifest in the structure, we took representative snapshots of the structures in each case from MDS (Figure 5). We find that the structural differences highlighted by SASA values are reflected in the conformational changes of the RNA molecule. In the WT structure, exon 11 exhibits a relatively compact conformation with limited exposure to the solvent. This suggests that the WT sequence of the RNA forms a stable secondary structure that may shield important binding sites from the surrounding environment. One example includes the 5’SS, which maintains a relatively closed structure in the WT pre-mRNA. Upon MBNL1 binding, presumably, the 5’SS and 3’SS become more accessible for U1 and U2 snRNP binding and defining exon 11 to promote the inclusion of this exon in the mRNA product.

**Figure 5.**
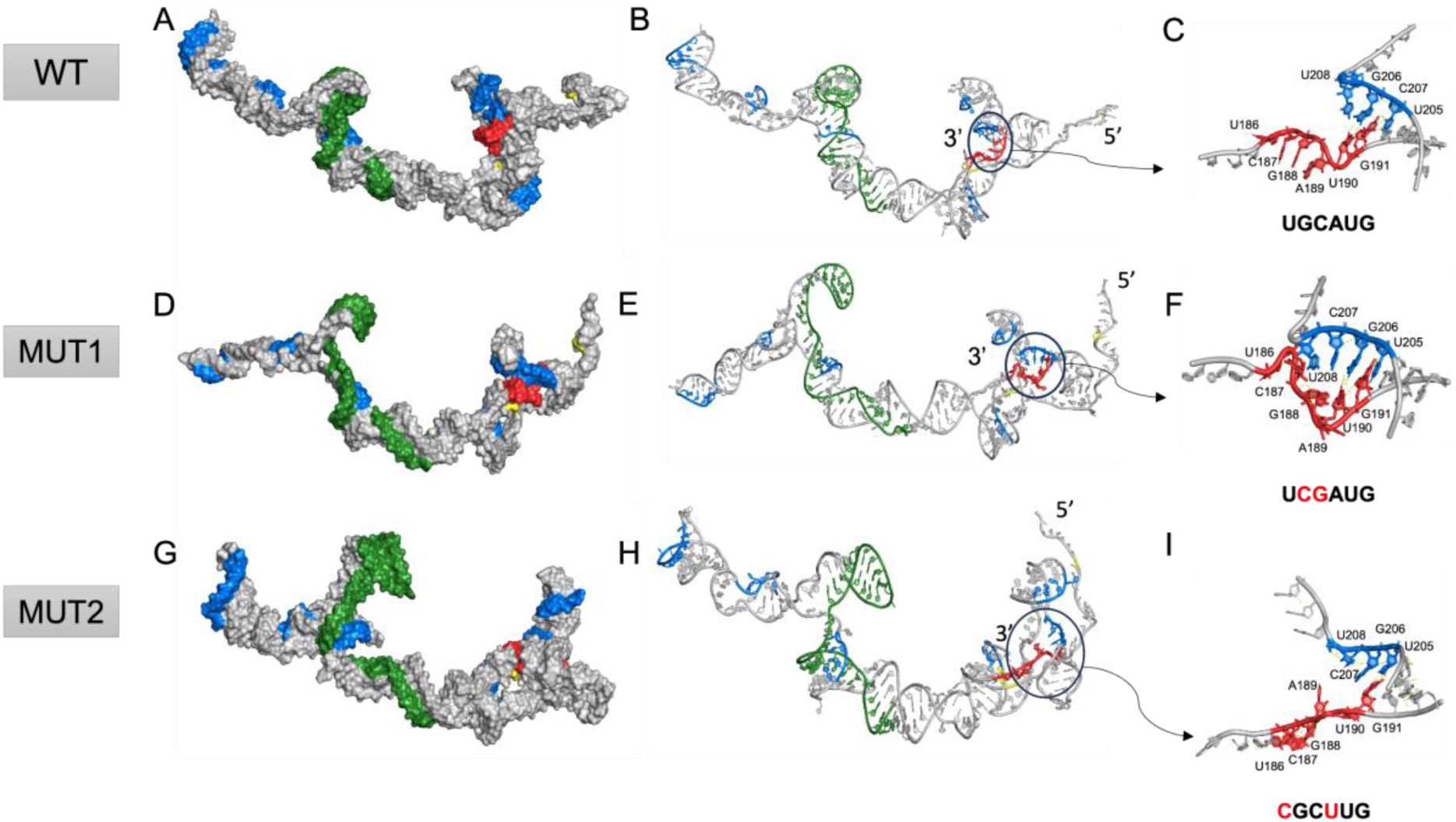
The figure showcases the surface (**A, D, G**) and cartoon representations (**B, E, H**) of the WT, MUT1, and MUT2 RNA structures, with a zoomed-in view (**C, F, I**) of the R-BS. The surface representation provides a visual depiction of the SASA of the RNAs, highlighting potential differences in their exposed regions due to the mutations. The cartoon representation offers insights into the overall secondary structure and conformational changes resulting from the introduced mutations. The zoomed-in view focuses on the R-BS, enabling a detailed examination of how MUT1 and MUT2 may impact the RNA’s interaction with RBFOX1.

In the MUT1 structure, exon 11 is more exposed to the solvent compared to WT. The increased SASA values align with the structural shift towards a more open conformation for this region of the *INSR* pre-mRNA. This more open exposure of exon 11 may affect the binding accessibility for regulatory proteins like MBNL1 and RBFOX1. The 5’SS showed a more solvent-exposed conformation, in agreement with the SASA analysis. The more exposed structure of this region of the pre-mRNA doesn’t correlate with strong MBNL1 splicing regulation, suggesting that the changes in RNA structure are caused by MUT2 blunt MBNL1 regulation. Interestingly, we observed that the MUT2 conformation of exon 11 and 5’ SS are closer to the WT pre-mRNA, with slightly less solvent exposure than MUT1. The shift of MUT2’s RNA accessibility (SASA scores more similar to WT) and snapshot structure similar to WT is consistent with the experimental results, with MBNL1 showing strong splicing regulation of both WT and MUT2.

In the snapshots of the structures, we observe the direct impact of the mutations on the R-BS. The R-BS is single-stranded in the WT structure (zoom in on R-BS in Figure 5C), while in contrast, the R-BS is in a double-stranded conformation with an adjacent MBNL1 recognition site in the MUT1 structure (Figure 5F). In the MUT2 structure, the R-BS is not base-paired and adopts a similar conformation to that observed in the WT structure (Figure 5I). The collective findings from both the SASA analysis and the representative snapshots provide a detailed insight into how the mutations influence the RNA’s structural dynamics compared to the WT. To see more details, Figure S1 in the supplementary material presents three short simulation videos for (A) WT, (B) MUT1, and (C) MUT2.

In addition to solvent accessibility, the structural stability of the binding sites on the RNA can contribute to protein binding. To explore the stability of the *INSR* pre-mRNA fragment, we used root mean square fluctuations (RMSF) as a metric to assess nucleotide fluctuation, with lower RMSFs correlating with higher stability and higher RMSFs correlating with lower stability. As shown in (Figure 6), we found that the fluctuation patterns of the nucleotides for MUT1 are remarkably different compared to MUT2 and WT. While MUT2 and WT appear to be primarily stable (low RMSFs throughout), MUT1 is characterized by larger fluctuations, particularly at the 3’SS and nearby MBNL1 binding site, M-BS4. These folding results suggest that WT and MUT2 adopt similar structures that are stable, while MUT1 samples more structures, which may not be favourable for MBNL1 binding and splicing regulation.

**Figure 6.**
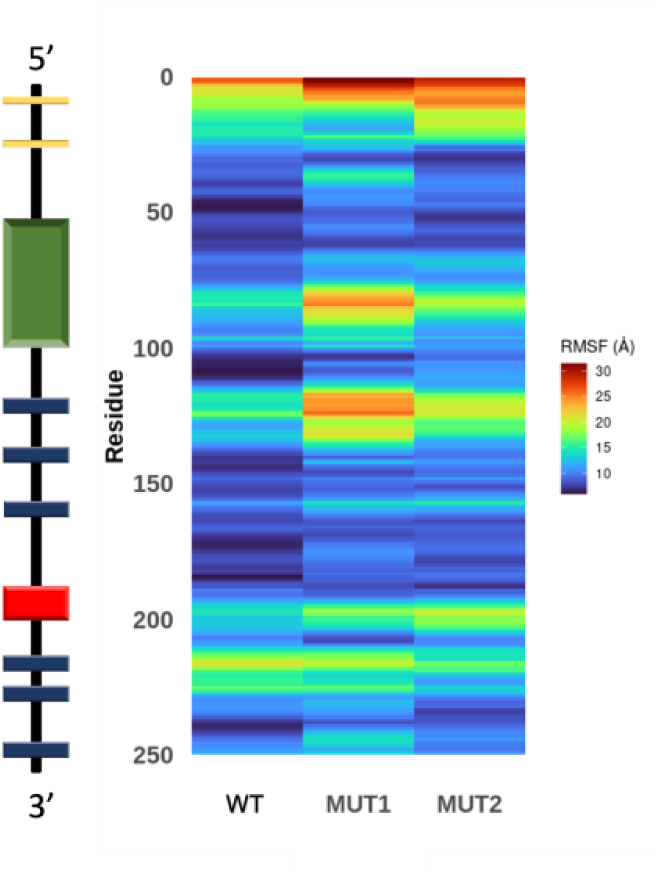
The RMSF heat map color scale indicates the magnitude of the RMSF values.The color gradient ranges from dark blue, representing lower RMSF values, to red, representing higher RMSF values. The ischemic region on the left side of the heatmap represents various segments of the modelled INSR pre-mRNA, as illustrated in Figure 1 earlier.

### Improving splicing with an antisense oligonucleotide that blocks predicted RNA structure element

To test the functionality of the predicted RNA structure of the *INSR* pre-mRNA, we used ASOs, which have been used extensively to modulate pre-mRNA splicing (60). For these studies, the endogenous *INSR* pre-mRNA was studied in the tetracycline-inducible HA-MBNL1 HEK-293 cell line used for the minigene reporter studies. Two positive control ASOs targeting the 3’SS (3’SS ASO, Figure 7A, B) and 5’SS (5’SS ASO, Figure 7A, B) were designed to block exon 11 inclusion. As expected, these ASOs inhibited the inclusion of exon 11 in the absence and presence of MBNL1, although a modest increase of exon 11 was observed when MBNL1 was added, and 5’SS and 3’SS blocking ASOs were present (Figure 7C). As previously observed (50), the deltaPSI for the endogenous *INSR* exon 11 was larger compared to minigene (0.65 Figure 7C vs 0.31 Figure 2A). The large change in PSI and complete sequence of the *INSR* pre-mRNA makes this substrate a good candidate for testing the role of a predicted RNA regulatory element.

**Figure 7.**
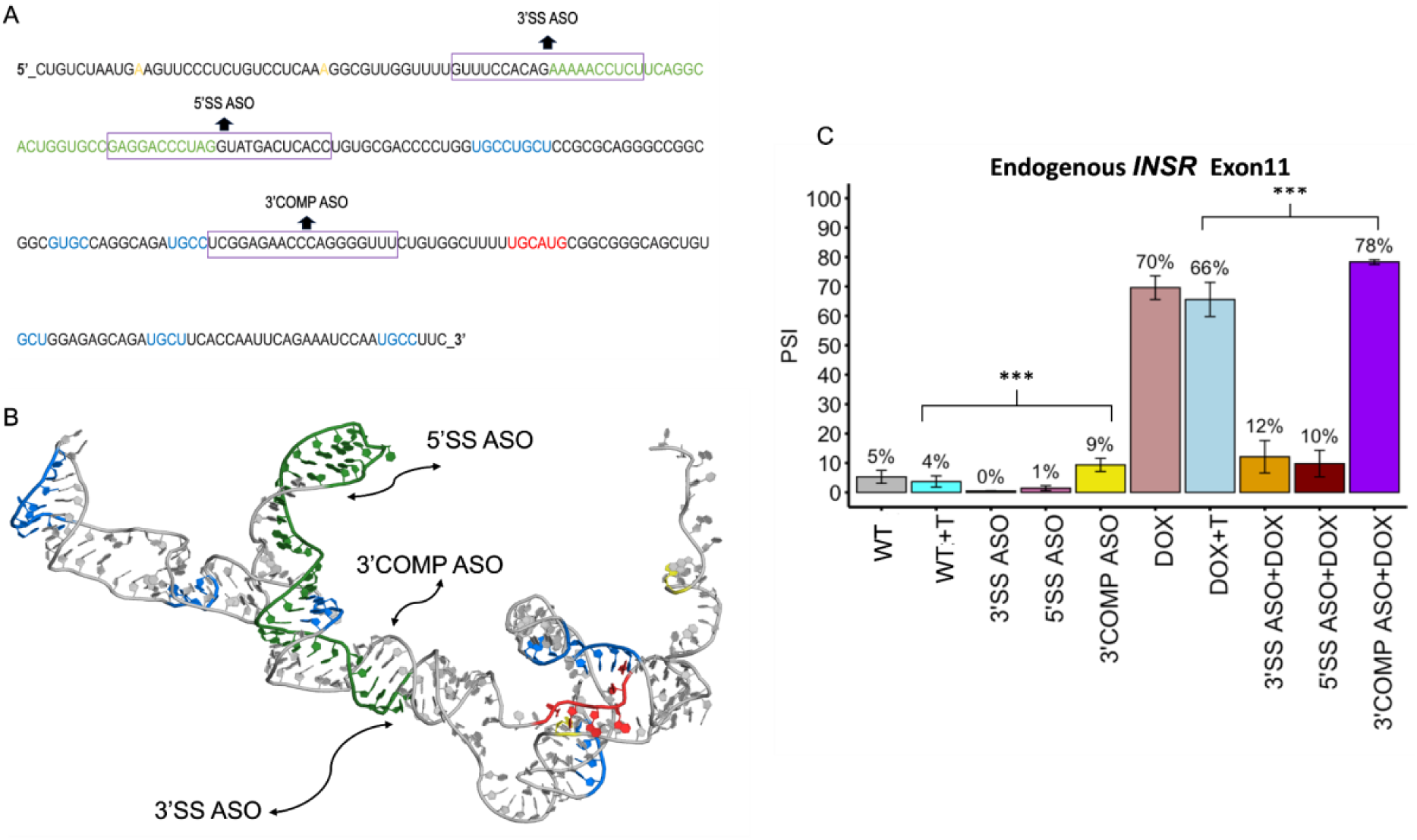
The primary sequence is shown in (**A**), with three sites targeted with ASOs (3’SS, 5’SS, and 3’COMP). **(B)** The modelled tertiary structure of *INSR* pre-mRNA with binding sites for ASOs marked with arrows. **(C)** The bar chart illustrates the PSI values of endogenous *INSR* exon 11 under various treatment conditions. Each treatment group is listed on the x-axis (T = transfection reagent, Dox = Doxycycline plus ASO treatment). Error bars represent standard deviations. Asterisks atop lines connecting treatment groups denote significance (p-value < 0.0001 (***)).

Guided by insights from our computational model and structural analysis, we predicted that disrupting the double-stranded structure at the 3’SS (Figure 7A) could facilitate greater accessibility for RNA-binding proteins (RBPs) and the splicing machinery. An ASO (3’COMP) that is designed to bind the 3’SS (Figure 7B) was tested for modulating splicing. In the absence of MBNL1 induction, the inclusion of exon 11 increased by 5% (4% to 9% with a p-value < 0.0001, Figure 7C). In the presence of MBNL1, this impact of the 3’COMP ASO was greater, with a 12% change (66% to 78%), demonstrating that this ASO positively modulates exon 11 inclusion at low and high MBNL1 levels.

## Conclusion

We used a combination of computational and experimental approaches to begin to unravel the complex regulatory mechanisms underlying the splicing of *INSR* exon 11. We demonstrated that the RBFOX UGCAUG motif within *INSR* intron 11 is important for the co-regulation of exon 11 inclusion by MBNL1 and RBFOX1. Mutating the UGCAUG motif to a <u>C</u>GC<u>U</u>UG motif containing an optimal YGCY MBNL1 motif partially restored MBNL1-mediated splicing regulation, suggesting MBNL1 binding this region of the pre-mRNA is important. We have also found that these modest two nucleotide changes (MUT1 and MUT2) significantly impact the predicted structure of the pre-mRNA.

Our computational simulations provide structural insights into the impact of mutations that alter the pre-mRNA conformation on the expected splicing outcome. Our simulations highlight a crucial distinction, predicting a mismatch bulge for MUT1, while MUT2 and the WT exhibit exposed single-stranded RNA regions. This mismatch bulge emerges as a key mechanism, underscoring the significance of our findings in elucidating the intricate regulatory dynamics of INSR exon 11 splicing. The SASA analysis revealed changes in solvent accessibility, with MUT1 exhibiting increased exposure of motifs known to be important for splicing regulation. The RNA structural changes corresponded to the functional splicing outcomes, indicating the interplay between RNA structure, protein binding, and splicing regulation. Overall, our integrative approach provides insight into the intricate mechanisms governing *INSR* exon 11 splicing regulation, highlighting the importance of both RNA sequence and structural determinants in regulating AS. Future studies could explore the dynamic interactions between pre-mRNA and RNA-binding proteins to deepen our understanding of splicing regulation in health and disease.

## Acknowledgments

Special thanks to the members of Berglund, Chen and Reddy labs for helpful discussions and experiment advice. We thank our RNA Institute colleagues for the general support and guidance.

