## Supplemental videos for "Mutation of two intronic nucleotides alters RNA structure and dynamics inhibiting MBNL1 and RBFOX1 regulated splicing of the Insulin Receptor": manuscript-vids (1).docx

**Supplemental information: Mutation of two intronic nucleotides sufficiently alters RNA structure and dynamics to inhibit regulated splicing of the Insulin receptor**

Zohreh R. Nowzari^1,2^, Melissa Hale^3^, Joseph Ellis^2,4^, Samantha Biaesch^1^, Sweta Vangaveti^2^, Kaalak Reddy^2,5^, Alan A. Chen^1,2*^ and J. Andrew Berglund^2,5*^

Figure S1.

A

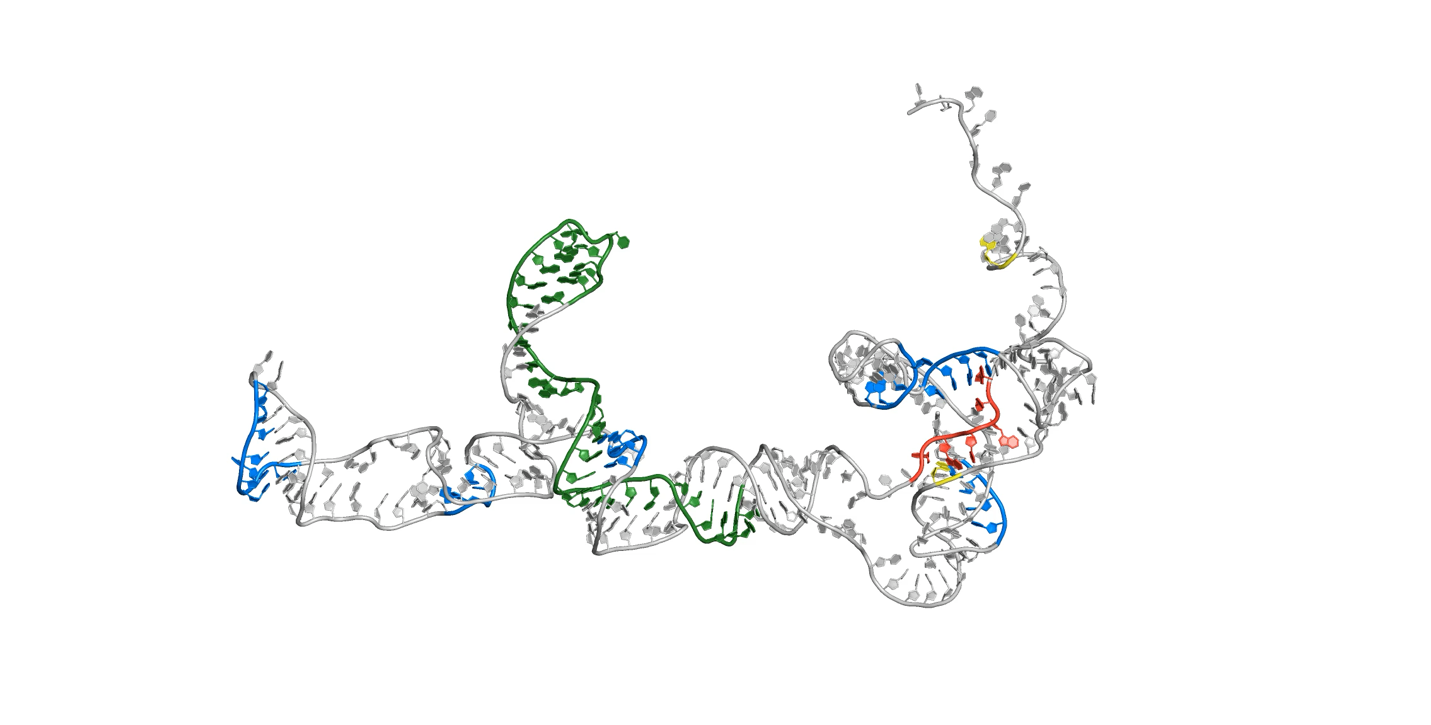

B

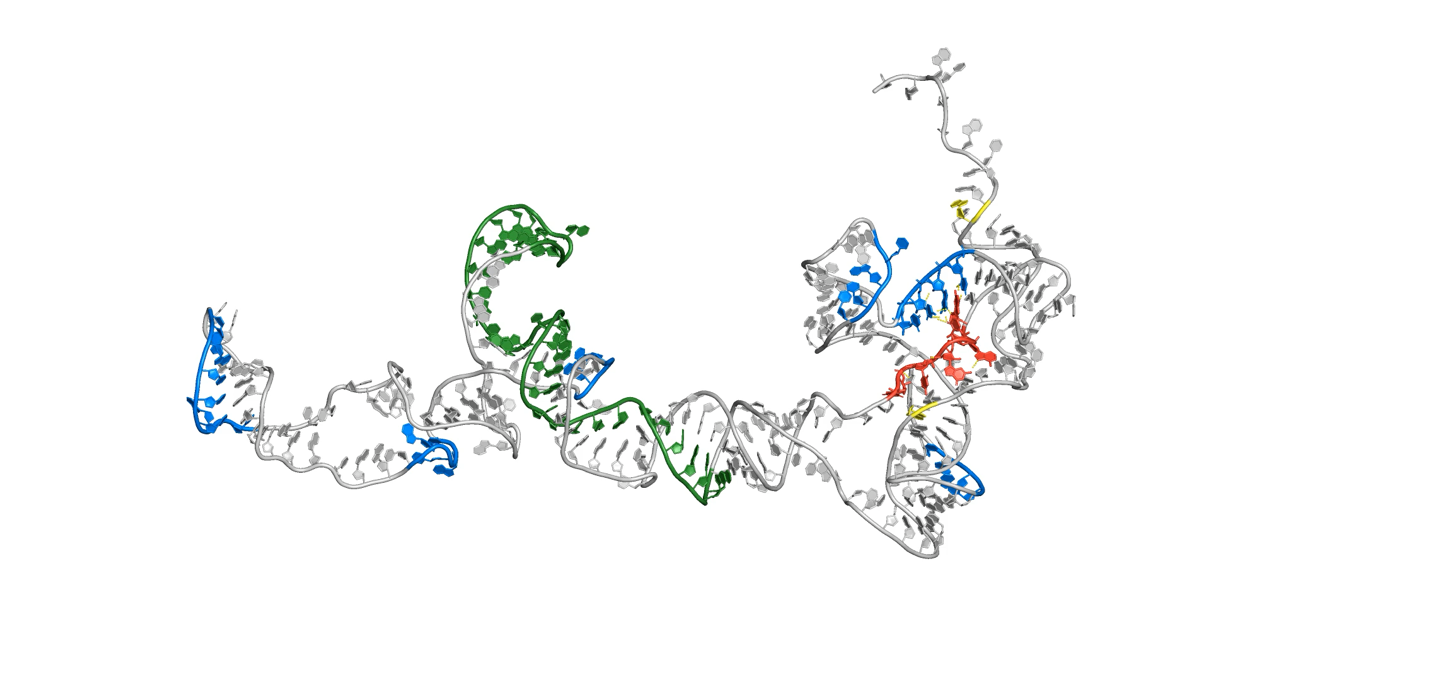

C

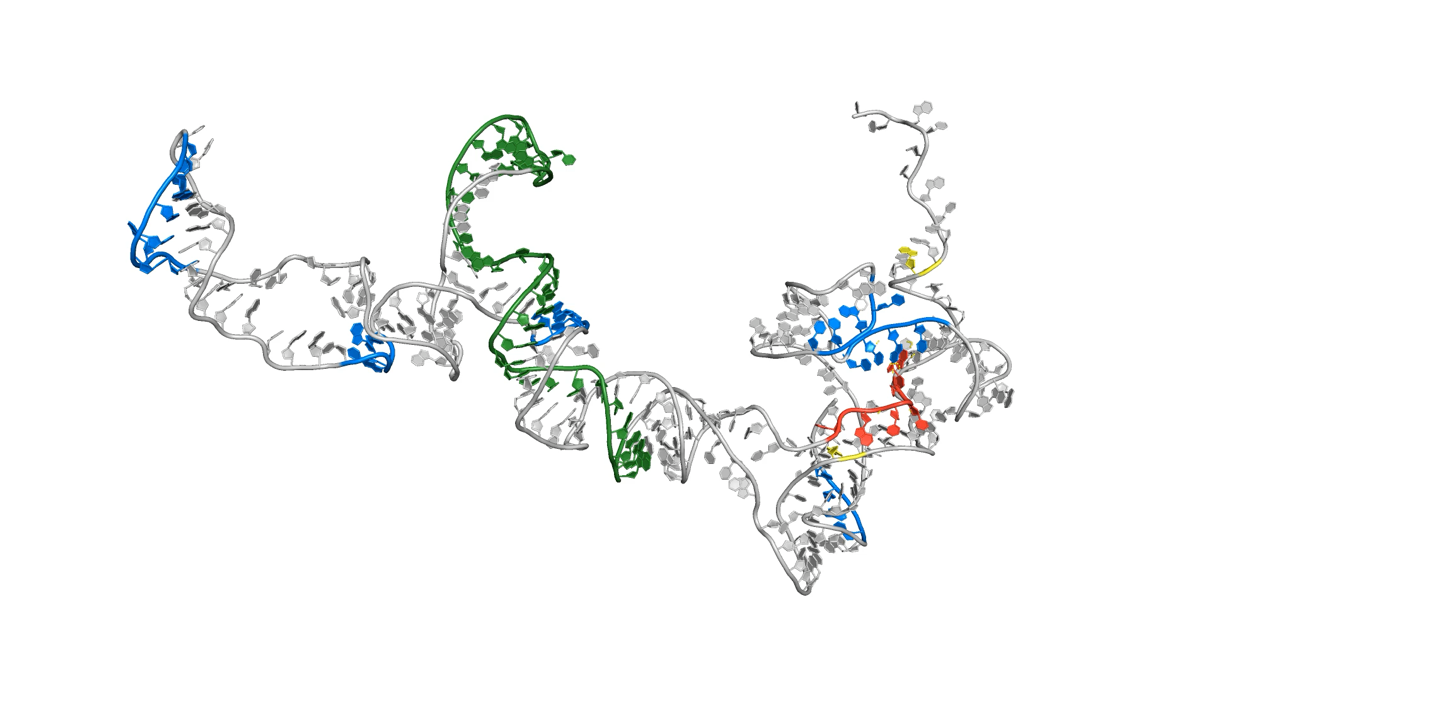

**Branch points (BP1,2) =Yellow**

**MBNL1 Recognition Sites(M-BS) = Blue**

**Rbfox1 Recognition Site(R-BS) = Red**

**Exon11(Ex11) =Green**

The figure illustrates three 90 ns MD simulation videos, each starting with different structures: **(A)** WT, **(B)** MUT1, and **(C)** MUT2.
